# Evaluating Transcriptomic Integration for Cyanobacterial Constraint-based Metabolic Modelling

**DOI:** 10.1101/2025.02.10.637384

**Authors:** Thomas Pugsley, Guy Hanke, Christopher Daniel Patrick Duffy

## Abstract

Metabolic modelling has wide-ranging applications, including for the production of high-value compounds, understanding complex disease and analysing community interactions. Integrating transcriptomic data with genome-scale metabolic models is crucial for deepening our understanding of complex biological systems, as it enables the development of models tailored to specific conditions, such as particular tissues, environments, or experimental setups. Relatively little attention has been given to the assessment of such integration methods in predicting intracellular fluxes. While a few validation studies offer some insights, their scope remains limited, particularly for organisms like cyanobacteria, for which little metabolic flux data are available. Cyanobacteria hold significant biotechnological potential due to their ability to synthesize a wide range of high-value compounds with minimal resource inputs [21]. The impact of specific methodological decisions on integration, however, has scarcely been assessed beyond human models, with no thorough exploration of parameter choices in valve-based integration methods. By implementing a novel analysis pipeline, we evaluated these methodological decisions using the genome-scale model for *Synechocystis* sp. PCC 6803 (iSynCJ816 [17]) with existing transcriptomic data in biomass-optimised scenarios. Our analyses indicate that selecting an appropriate integration method may not always be straightforward and depends on the initial model configuration - a factor which is often overlooked during integration. By evaluating sets of methods, we identified a trade-off between the buffering of light into the system and maintenance of flux near system boundaries. We suggest that the use of the lazy-step mapping function with importance-based scaling results in the best predictions, particularly when these can be validated with experimental data. When using one-size-fits-all scaling with the lazy-step mapping function, it appears preferable to use light buffering to avoid inappropriate bound changes near the photosystems, a factor which importance-based scaling may help to compensate for. In cases where no experimental data can be used for validation, the novel thresholding approach could be adopted as this showed some improvements upon the standard Lazy method.

## 2 Introduction

Constraint-based metabolic models are mathematical representations of cells which enable system-level analyses of metabolism. Metabolic models are useful tools for the prediction of metabolic fluxes and cellular phenotypes, with wide ranging applications across biotechnology. [41]. Covert et al. (2001) published the first method to integrate transcriptomic data with metabolic models, enabling model simulations to account for gene regulation [8]. In the following decades, many more integration methods were developed, allowing researchers to capture condition-specific properties of metabolic flux distributions, enhancing the specificity of downstream analyses [6] [47] [7] [2] [1].

Integration methods can be broadly divided into two categories: switch-based and valve-based [16] [37]. Switch-based methods use thresholding to categorise reactions based on their predicted activity, with reactions of low predicted flux being dropped from the metabolic model. This type of integration has predominantly been used to model human cell types, with previous studies having examined the impact of specific methodological decisions during integration [28] [9] [18]. Richelle et al. (2019) examined gene mapping types, thresholding and the order of those steps for the creation of human tissue-specific models while Gopalakrishnan et al. (2022) mainly focused on algorithmic details and changing thresholding cut-offs in *E. coli*, Chinese Hamster Ovary (CHO) and a renal cancer cell line. Joshi et al. (2020) also examined transcriptomic integration with a cancer cell line model, focused mainly on the influence of thresholding values. Valve-based methods involve modifying reaction bounds in a continuous manner, with bounds being relaxed for up-regulated reactions and tightly constrained for down-regulated reactions. Although enzyme activity is not always directly correlated with transcript levels, these methods assume that gene expression can be used as a ‘soft’ constraint to approximate an upper bound for reaction rates. It has been argued that valve-based approaches are preferable as they do not suffer from the loss of fine-grained expression changes in the same way as switch-based methods [22].

METRADE [2] and E-flux [7] are examples of valve-based integration methods, both of which have been applied to microbial metabolic models. In the case of E-flux, Colijn et al. (2009) used their newly developed integration method to model mycolic acid biosynthesis, predicting the responses of drugs. Since then, it has become a popular integration method and a second implementation has been developed, E-flux2 [20]. METRADE, on the other hand, developed by Angione and Lió (2015), has been implemented as a multi-objective optimisation problem. Originally utilised for the creation of multi-omic models of *E. coli*, METRADE has since been applied to the cyanobacterial species Synechococcus sp. PCC 7002. Using a hybrid machine learning FBA pipeline, repsonse mechanisms to light and salinity fluctuations were detected - a finding which was not possible from the analysis of the transcriptomic data alone [38].

As pointed out by Machado and Herrgård (2014), publications which introduce novel integration methods often illustrate their applications with specific case studies, resulting in little uniformity within the literature. For example, Colijn et al. (2009) reported reassuring results that E-flux was able to correctly predict 7 out of 8 known fatty acid inhibitors as well as their specificity for biosynthesis. In the case of METRADE, Angione and Lió (2016) validated their integration method using a series of 14 protein expression profiles with paired growth rates, yielding strong correlations between predicted and experimental data. In both cases, while some form of validation has taken place, neither approach allows for the direct comparison of different implementations, or any deeper methodological evaluation, such as scaling strategy or parameter selection. Although previous studies have been motivated by the need to benchmark integration methods using common datasets [35], such studies remain relatively uncommon, partly due to the difficulty of obtaining sufficient paired fluxomic and transcriptomic data [22].

13C metabolic flux analysis (13C-MFA), in which cells are fed 13C-labelled substrates and enrichment patterns reconstructed, is widely accepted as the gold-standard for quantifying flux through central carbon metabolism [40] [45] [4]. For this reason, 13C-MFA measurements are typically used to validate flux predictions by metabolic models and the use of integration methods, as they provide systems-wide datasets for comparison [24]. However, 13C-MFA data is notoriously challenging to produce and requires highly specialised expertise [20]. For this reason, few extensive 13C-MFA datasets exist (relative to transcriptomic studies), even for well-studied organisms such as *E. coli* and *S. cerevisiae* [10]. Unsurprisingly, there is even greater dataset scarcity among cyanobacteria, with only 3 central carbon flux distributions having ever been published alongside paired transcriptomics [5] [43] [42] [44]. Methods for assessing flux predictions from context-specific models in cyanobacteria are therefore severely limited. Complicating matters further, cyanobacterial systems present distinct challenges compared to their heterotrophic counterparts. In particular, the inference of photosynthetic behavior from transcript profiles has yet to be robustly assessed, and optimal lighting configurations remain relatively unexplored.

In this study, we present a novel pipeline to evaluate the performance of integration methods for the creation of context-specific models of *Synechocystis* sp. PCC 6803 (hereon referred to as *Synechocystis*) using existing expression profiles in CyanoExpress [13]. By creating relative growth rate traces using optical density (OD) data, we derived 7 datasets for validation. The use of curve fitting allowed for growth rate estimation in cases where authors did not measure OD at the time of RNA harvest, thereby extending the number of viable datasets suitable for downstream analysis. Since it is common for publications to analyse the onset of a transcriptional responses in time series, we were able to capitalise on having multiple observation prediction targets measured by the same set of authors. Using Flux Balance Analysis (FBA) [27], we predicted growth rates at relevant time points. Early on, after the onset of an environmental stress, it is typical to expect smaller global transcript fold changes (relative to a hypothetical control) than those from the later adaptive phases. Typically, lower scaling values are required during mapping for high global fold change transcripts to impact biomass predictions than those with low global fold changes. By analysing growth rate trace similarity over the course of stress onset, we could more intricately assess scaling parameter performance and strategy as they relate to the entire cellular response. We assessed growth rate trace similarity using Dynamic Time Warping (DTW) [32] so that distance measures would not heavily penalise similarity scores if traces were correctly shaped, but temporally misaligned. This feature was particularly important given there may have been differences in how authors disclosed their RNA harvesting times. Finally, to account for differences in sample size, we generated dataset-specific null distributions and used resulting p-values to infer predictive performance. By using biomass predictions as a proxy for system behaviour, we were able to investigate the impact of methodological decisions at different stages of integration and suggest improvements for more robust future analyses.

## 3 Methods

### 3.1 Transcriptomic data

Expression profiles deposited on CyanoExpress 2.3 (pre-processing and normalisation details available from: http://cyanoexpress.sysbiolab.eu/) were screened for their suitability for downstream analysis. Each transcriptomic dataset was required to have expression profiles from a minimum of 3 independent timepoints and have paired OD data to infer growth rates from (within their associated source papers). WebPlotDigitizer was used to extract experimental data (Supplementary figure 3) from published OD plots [30]. Transcriptomic log-fold change values were converted to fold-change (so that wild-type expression was equal to 1) before use for metabolic modelling.

### 3.2 Growth Rate Derivation

Since absolute growth rate determination is not possible from optical density data alone, we derived relative growth rates for validation. These relative growth rate traces were derived by inferring values from fitted OD curves. For each OD plot, either an exponential, exponential rise, or logistic curve was fit with the resulting curve’s parameters used to estimate the growth rate as outlined below.

We consider OD to be

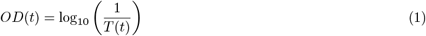

where *T* (*t*) is the *transmittance* of the sample as a function of time. In the limit of an infinitely far detector, the transmittance would be given by

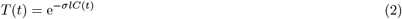

where *σ* is the scattering cross-section of the particles, *l* is the optical path length, and *C*(*t*) is the time-dependent concentration. Without knowing *σ* and *l*, a *relative concentration* may be derived:

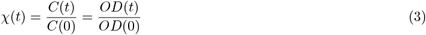

where *C*(0) and *OD*(0) are the concentration and OD at *t* = 0. We therefore define the *relative growth rate*:

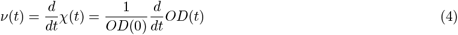

#### 3.2.1 Exponential Fitting

If the experimental OD curve shows exponential growth, we assume:

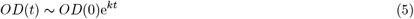

where *k* is a phenomenological rate constant. Substituting into Eqn. (4) gives:

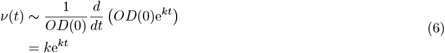

#### 3.2.2 Exponential Rise Fitting

If the experimental curve rises exponentially before plateauing, we use the basic functional form:

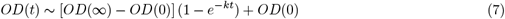

where *OD*(∞) is the *OD* level at the plateau. We calculate relative growth rate:

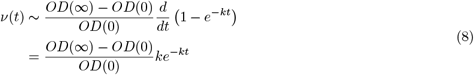

#### 3.2.3 Logistic Fitting

Finally, for sigmoidal OD curves, in which the initial lag phase and plateau are clearly discernible, we use the logistic function:

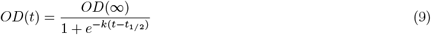

where *k* is the *logistic growth rate constant* and *t*_1*/*2_ is the time at which the OD reaches *OD*(∞)*/*2. We calculate relative growth rate as:

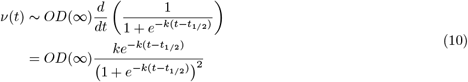

### 3.3 Optimisation

Using the COBRA Toolbox version 3 [12] for MATLAB, Flux Balance Analysis (FBA) was used to predict growth rates at different time points using the *Synechocystis* metabolic model, iSynCJ816 [17].

In metabolic modelling, the metabolic network is represented as a stoichiometric matrix **S**, where each element *S*_*ij*_ corresponds to the stoichiometric coefficient of metabolite *i* in reaction *j*. The matrix **S** has dimensions *m* × *n*, where *m* is the number of metabolites and *n* is the number of reactions.

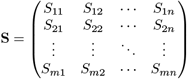

The system is assumed to be at steady-state, resulting in this system of linear equations:

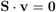

where **v** is the vector of reaction fluxes. Each element *v*_*j*_ in **v** represents the flux through reaction *j*. For all simulations, the objective function was set to the autotrophic biomass equation.

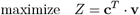

where **c** is a vector that specifies the objective function. Reaction fluxes are subject to upper and lower bound constraints. Ultimately, these are determined by each integration method.

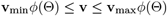

where **v**_min_ and **v**_max_ are vectors of the default minimum and maximum allowable fluxes for each reaction and function *ϕ*(*θ*) is the mapping function used for integration. The FBA problem is formulated as a linear programming (LP) problem and solved using the MATLAB GLPK solver:

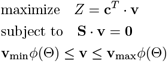

### 3.4 Starting Model Configurations

In the context of this study, model ‘configurations’ refer to the initial bound setup of the metabolic model prior to bound alteration by an integration method. We tested integration methods starting with two different model configurations. For both configurations, all non-zero bounded reactions in the default model were set to an arbitrary value of 10 or -10. In configuration ‘Baseline’, allowable flux through the photon input reaction was set to half of that of the rest of the system bounds. The purpose of this was to create a buffering effect so that the supply of light to the wider system was less likely to be influenced by transcriptional changes associated with the photosystems. For the other configuration, ‘Max’, photon import was capped at the same upper bound as the rest of the system.

To simulate autotrophic growth, glucose import was switched off, bicarbonate import was modelled as the sole source of carbon to the system and the autotrophic biomass reaction was maximised.

### 3.5 Integration Methods

Key features of each integration method are shown in Table 1.

**Table 1:** Key features of integration methods.

| Name | Mapping function | Thresholding | Scaling |
| --- | --- | --- | --- |
| Lazy | Lazy-step | No | One-size-fits-all |
| Lazy threshold | Lazy-step | Yes | One-size-fits-all |
| Lazy importance | Lazy-step | No | Reaction-specific |
| Linear | Linear | No | One-size-fits-all |

#### 3.5.1 Gene expression to reaction bound conversion

Mapping functions are used to map ‘reaction expression’ values to bound changes. In order to estimate reaction expression from gene expression data, it is required to consult the boolean expressions in the gene-reaction rules of iSynCJ816. If only a single gene influences a given reaction’s flux, its expression value is determined to be the reaction expression value. To model multi-gene enzymes or isozymes, we used the approach from METRADE [2]. If *x*_*ij*_, *i* = 1, …, *p* are the gene expression values in a dataset with genes *s*_*j*_, *i* = 1, …, *p*:

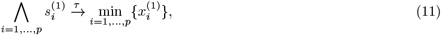

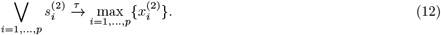

where Λ *s*^(1)^ is a gene set contributing to an enzymatic complex and Λ s^(2)^ is a gene set for an isozyme.

#### 3.5.2 Mapping function

Once reaction expressions have been derived for all reactions in iSynCJ816, they can be converted into new reaction bounds using a mapping function (as above):

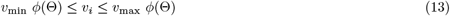

In this report, two mapping functions were used: the lazy-step function (as used in METRADE) [2]

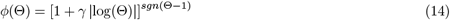

and a linear mapping function, similar to the approach used in the popular valve-based method E-flux [7]:

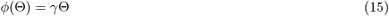

where Θ is the computed reaction expression.

It is important to consider that these integration methods are not explicit implementations of METRADE or E-flux, but are closely related. The approach taken in this report facilitates direct comparison of specific methodological interventions, rather than a rigid assessment of METRADE or E-flux’s performance.

#### 3.5.3 Thresholding

Thresholding is a feature more commonly used in conjunction with switch-based integration to determine the activity/inactivity of model reactions and has been relatively unexplored in the context of valve-based integration. We created novel integration methods using thresholding in which bound changes were only applied if reaction expressions were at least 10% above or below the control. For methods that did not use thresholding, all reaction changes were converted into bound modifications. While the 10% value can be considered somewhat arbitrary, it is loosely based on expression discretisation advice from other integration methods like iMAT. Zure et al. (2010) suggest a threshold of half a standard deviation from the mean. [47] Applying this logic to our dataset results in an upper and lower threshold of approximately 10%, justifying our strategy.

#### 3.5.4 Scaling

In this report, we used two scaling strategies: ‘one-size-fits-all’ (in which *γ* in equations (14) and (15) was scaled to the same value across all reactions) and ‘reaction-specific’ - also referred to as ‘importance’. We define the importance of a gene in the same way as Angione et al. (2015) [2]. The gamma parameters tested, in the context of importance, relate to the maximum gene importance value. For each gene in each condition, gene variance values were computed from the expression profiles. The maximum gene importance value was used to normalise these variances such that

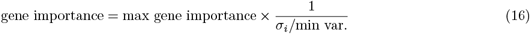

where *σ*_*i*_ is the variance of gene *i* and min var. is the minimum variance value of the variances dataset. For each gene in the model, we determined whether it was present in the expression profiles. Letting *G* be the set of genes in the model, and *D* be the set of genes in the transcriptomic dataset, gene importance *γ*_*i*_ is defined as:

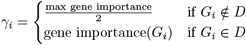

### 3.6 Assessment of predictions

#### 3.6.1 Data Normalisation

To allow meaningful comparisons between predicted and experimentally-derived growth rate traces, both datasets were scaled to a 0-1 range. Given a set of reference data 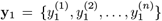 and prediction data 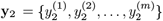, min-max normalisation for each data point 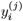 was performed using:

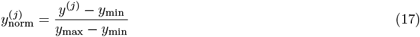

where *y*_min_ and *y*_max_ are the minimum and maximum values of the dataset, respectively.

#### 3.6.2 Dynamic Time Warping

DTW was used to assess temporal growth rate trace similarity. Given a reference **y**_1_ and prediction trace **y**_2_, for each pair of points (*i, j*), the squared difference (cost) between the elements of the sequences was computed as follows:

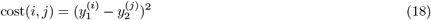

The DTW matrix was populated using the following recurrence relation:

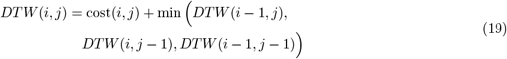

where *i* ∈ {1, 2, …, *n*} and *j* ∈ {1, 2, …, *m*}. The final DTW distance between the two traces is then given by:

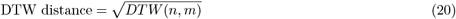

The final distance gives the optimal alignment that minimises the cumulative distance between both growth rate traces.

#### 3.6.3 P-value derivation

The number of observations in each dataset-specific reference trace can vary, necessitating the need for significance testing. For each set of predictions across all integration methods, traces were individually normalised to facilitate comparison using DTW. For each dataset, 100,000 random growth rate traces were computed based on the sample size of that dataset and subsequently max-min normalised. For each randomly generated trace, a DTW distance was computed by comparing to a given dataset’s experimentally-derived trace. These DTW distances formed a null distribution for each dataset, allowing for the computation of p-values as the proportion of randomly generated DTW distances lower than the prediction.

## 4 Results and Discussion

After screening the datasets in CyanoExpress, 7 sets of expression proiles were deemed suitable for downstream analysis (table 2). We implemented our novel pipeline using the 7 transcriptomic datasets, testing 1406 unique integration methods (fig. 1).

**Table 2:** Expression profile labels and their source publications - more details available from CyanoExpress [13].

| Plotting name | Description | Sample size | Reference |
| --- | --- | --- | --- |
| Cd | Cadmium stress | 9 | [15] |
| Blue light | Blue light growth | 6 | [33] |
| High light | High light stress | 6 | [34] |
| CrhR | CrhR-mutant; Low temperature | 3 | [26] |
| S starvation (H) | Sulphur starvation (no HEPES) | 3 | [46] |
| S starvation | Sulphur starvation | 7 | [46] |
| Iron stress | Iron stress | 5 | [14] |

**Figure 1:**
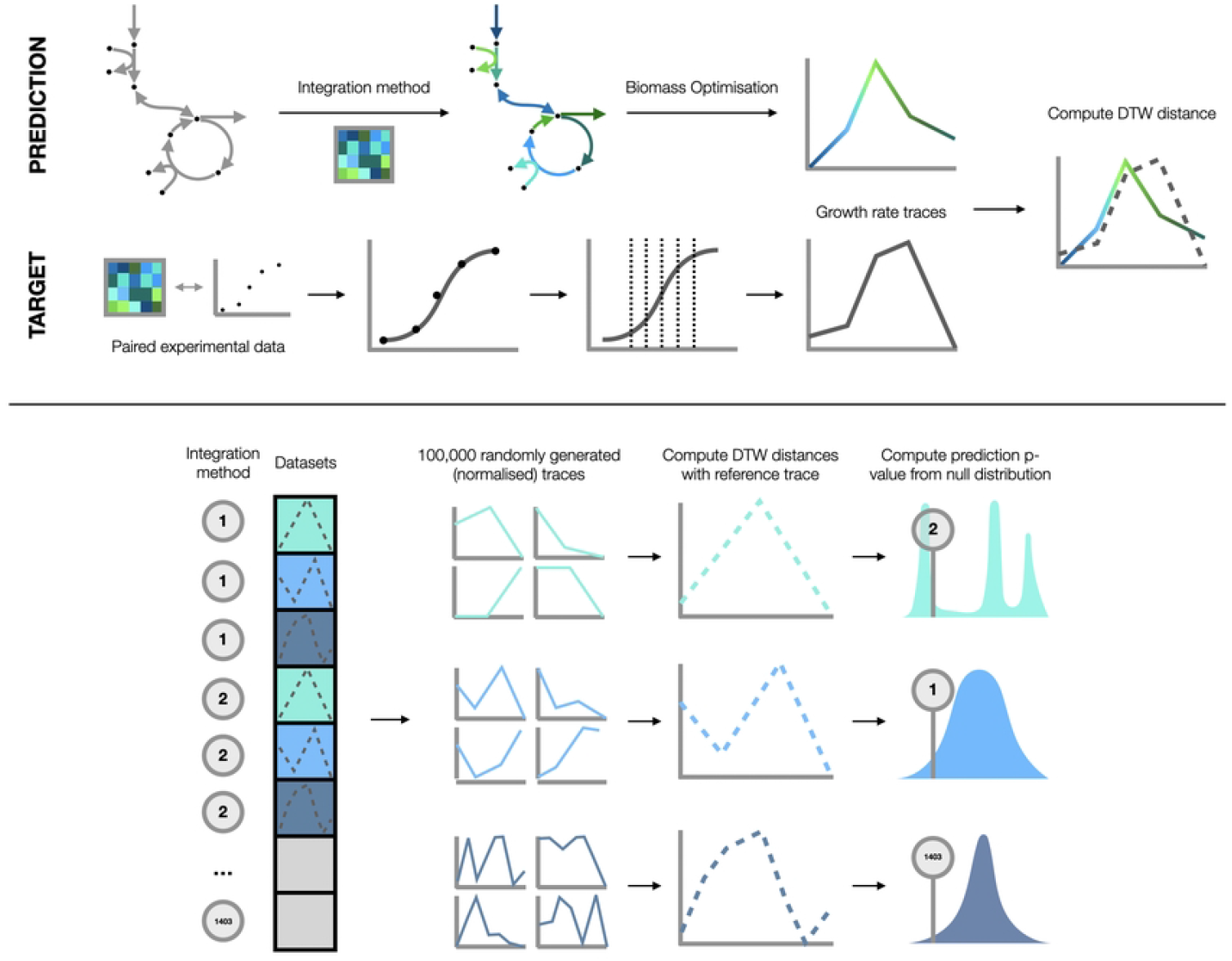
Methodology schematics: (Panel above) Optical density data associated with transcriptomic measurements at multiple timepoints were fit with curves, allowing for growth rate estimation in cases where authors did not measure OD at the time of RNA harvest. Curve fitting thereby extended the number of viable datasets suitable for downstream analysis. Integration methods were used to contextualise the metabolic model using the expression profiles from each timepoint. Maximising the biomass objective function was used to generate growth predictions at each time point, facilitating comparison of the predicted and experimentally-derived growth rate traces by computing DTW distances. (Panel below) Each integration method is tested on all time-series datasets - only 3 datasets are shown for simplicity (in reality, all 7 datasets were tested with 1403 unique integration methods). 100,000 random growth rate traces were computed for each set of datasets. Computing the DTW distances between these traces and the references, we created a series of condition-specific null distributions. P-values could then be determined to quantify the significance of predictive success. Across both panels, x axes represent time and y represent OD for curves and relative growth rates for lines. Axes are normalised time and normalised growth rate for line graphs.

### 4.0.1 Method robustness to scaling changes

When examining the scaling values which resulted in optimal predictions across different scaling strategies, it was clear that no lazy-step method was entirely robust to scaling parameter changes and that prediction performance dynamics were highly dependent on the condition being modelled (Supplementary figure 1). There was a reasonable level of consistency in optimal gamma values for the Lazy Threshold method, although there were only three viable parameter values to select from. An explanation for the infeasibility of the optimisation problem for gamma values *>*2 is not obvious. Use of the Linear mapping function with thresholding also resulted in infeasibility, although this occurred for all scaling values.

Comparing p-values for different gamma parameters within methods using the same mapping function allowed for a determination of the robustness of lazy-step function-based and linear-based integration methods specifically. For the gamma ranges tested, a broad level of consistency was observed for predictions using the Linear integration method for all conditions. In contrast, the analogous approach using the lazy-step mapping function, Lazy, showed a large degree of variability in prediction p-values (Supplementary figure 1). It was clear, however, that despite inconsistent prediction dynamics, upon selection of the optimal gamma parameter, across the board, the Lazy method could outperform the Linear method, for all conditions using the Baseline configuration, and all but the ‘Cd’ condition using the Max configuration in which optimal predictions were roughly on par (Supplementary figure 1). This is a simultaneous demonstration of the benefits of the Lazy-step function whilst highlighting the importance of scaling parameter selection.

### 4.0.2 Influence of model configuration

Regardless of whether ‘single condition-based scaling’ or ‘full dataset-based scaling’ was considered (scaling terms are defined in the legend of fig. 3), mean performance of individual methods across conditions also showed clear advantages for the use of the lazy-step function over linear mapping. Given the robustness of the linear mapping function to changes in gamma, it is unsurprising that this difference is less stark for full-dataset-based scaling comparisons (fig. 3f). There were also differences in mean predictive performance dependent on bound configuration. For the Lazy method, a buffered light input configuration resulted in better mean performance. It also appeared that 10% thresholding was able to relieve this dependence on configuration, and that the use of variance-based scaling (in the case of condition-based scaling), not only resulted in the lowest mean predictive performance, but also saw a shift in the effect of configuration choice relative to one-size-fits-all scaling. Our findings suggest that not only did importance-based scaling seem to compensate for a lack of light input buffering at the photosystems, but also resulted in an improvement in performance relative to the Baseline approach (fig. 3c,f).

**Figure 2:**
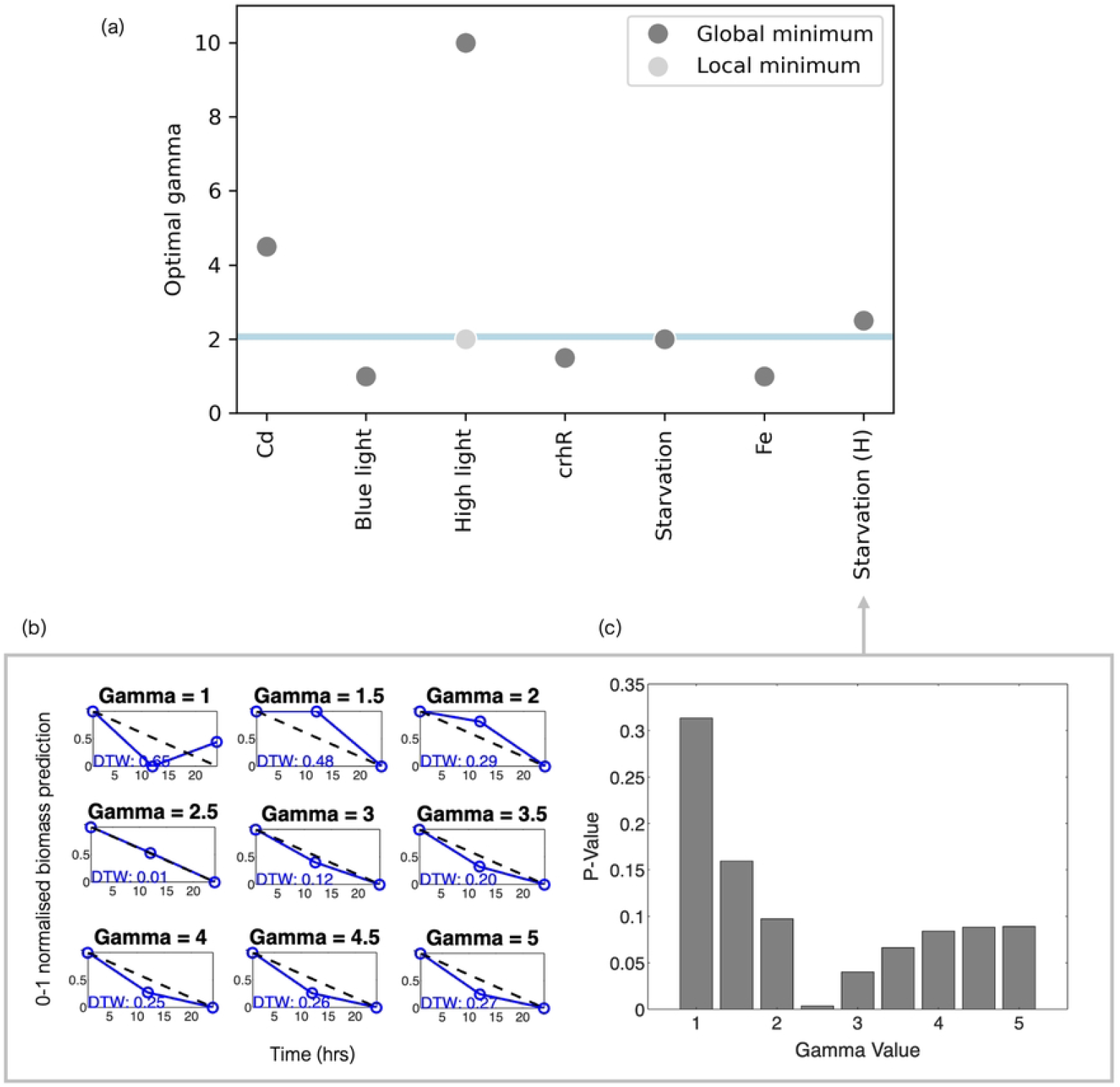
Optimal gamma values for the Baseline Lazy method across each set of conditions (a). In (a), we refer to a ‘global minimum’ which plots the optimal scaling value that yields the minimum p-value among those sampled. A ‘local minimum’ is also plotted for ‘High Light’ which indicates a visual dip (Supplementary figure 2) in prediction p-values close to the value (in magnitude) of the minimum p-value. The light blue line represents the mean optimal gamma value using the local minimum from ‘High Light’. A snapshot of the analysis pipeline to derive these optimal gamma values is also shown for ‘Starvation (H)’: the computation of DTW values (b) and resulting p-values (c).

**Figure 3:**
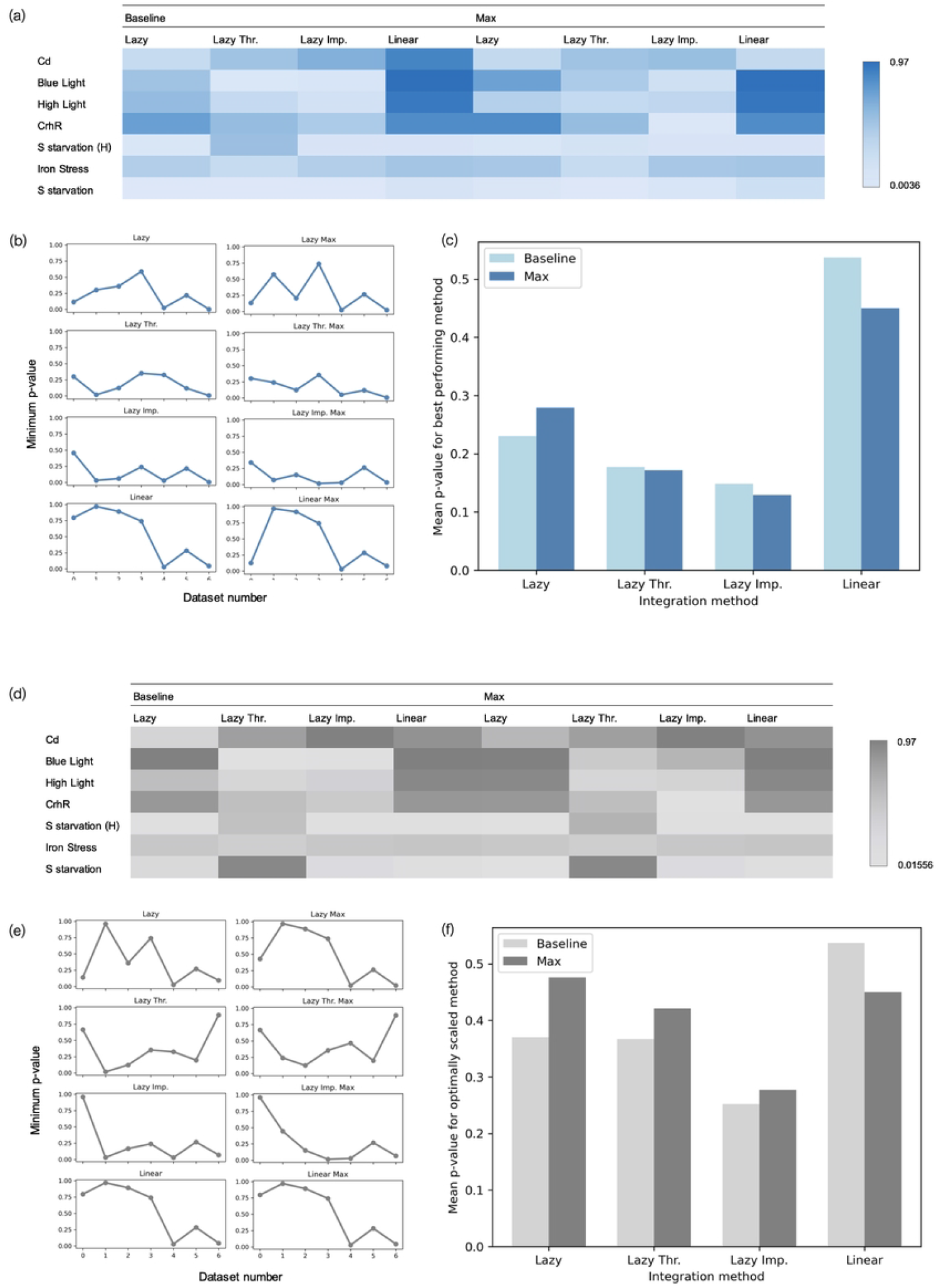
In blue (‘single condition-based scaling’): each method is optimally and individually scaled for every condition to demonstrate the potential for predictive success; In grey (‘full dataset-based scaling’): each method is optimally scaled across all condition datasets (lowest mean p-value after 0-1 dataset-specific normalisation) to show the performance of a specific method with a single scaling strategy. For each of these analysis types, heatmaps of p-values reveal the best performing predictions (a, d) alongside their associated line graphs (b, e) - datasets are displayed on the x axis and points are connected for ease of comparison, not to imply continuity. Bar charts display the mean p-value for each method type across all datasets (c, f) highlighting the overall performance of methods across all conditions.

The initial configuration of the model has the potential to alter predictive accuracy as the treatment of the model’s Boolean gene-reaction rules assume reaction fluxes are controlled by enzymes. Cyanobacterial photosystems, which are not themselves regulated like enzymes, are represented in iSynCJ816’s photosystem reactions, and are the first point at which transcriptional integration can have an effect on light entering the wider system. It can therefore be argued that these ‘points of entry’ for light should not be subject to strict bound changes based on expression profiles due to an inappropriate assumption that they are regulated in a manner similar to those of enzymatic reactions. To illustrate a violation of this assumption, a typical high light stress response can be considered (fig. 4a). It would be expected that as a long term strategy, photoystem proteins would be down-regulated to relieve the light load on downstream photosynthetic pathways. Without specifying explicit estimates for photon input in the metabolic model, after integration with transcriptomic data, such a response at the genetic level would result in a reduction of potential flux entering the system due to the shrinking of flux bounds at the photosystems. Such an assumption is inappropriate given the numerous non-transcriptional regulatory mechanisms which act to maintain flux through photosystem reactions [36] [29] [3]. Energy redistribution through phycobiliosome state transitions and dissipation of excess energy by non-photochemical quenching are key examples of this, both of which act with near immediacy [19] [31]. Therefore in our analyses, the ‘Baseline’ configuration was set up so that the upper bounds of the photosystem reactions (and all other reactions in the system) were twice as large as the available light, creating a buffer between any transcription-induced bound modification and light entering the system. In the ‘Max’ configuration, photosystem upper bounds were set to equal the available light, meaning any modification of the photosystem bounds could reduce light entering the system based on the gene-reaction rules.

**Figure 4:**
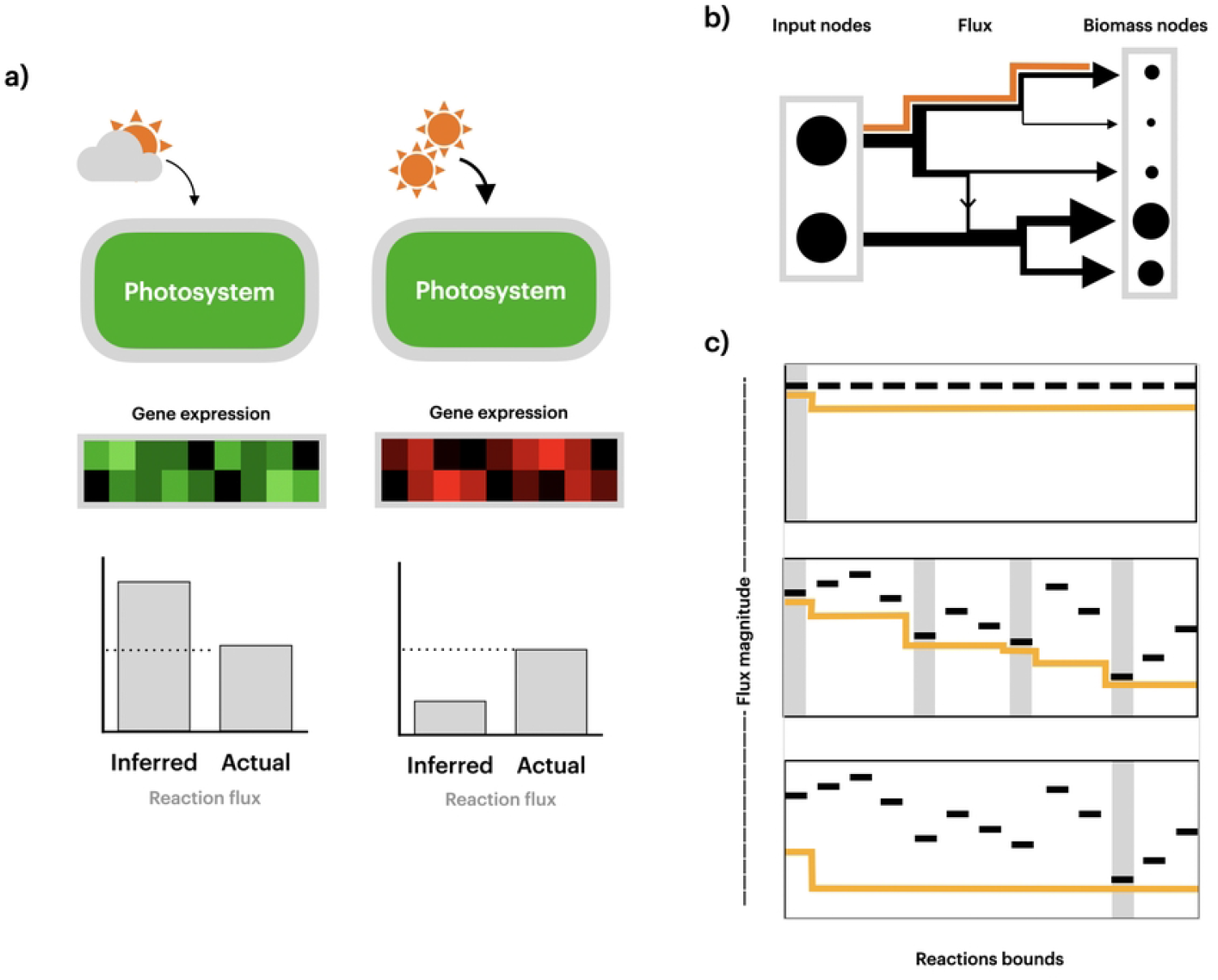
A schematic demonstrating the limitations of the gene-protein reaction rule assumption of the photosystems (A). (B) shows a hypothetical representation of flux dissipating from inputs to outputs. Arrow thickness represents flux magnitude and the orange trace follows a single pathway whose flux reduces downstream - the overall pattern of any pathway in the model. (C) shows 3 simplified cartoon representations of flux as it passes through reactions in the system: reaction upper bounds are represented as black dashed lines and flux magnitude is traced as a yellow line. Grey bars indicate the particular bounds which are responsible for shaping the solution of the pathway. The top panel shows the model bound configuration without any integration with transcriptomic data. The middle panel shows an example of the ‘Max’ configuration with integration where bounds are modified, flux is maintained near the reaction bound limits, and the photosystem reaction heavily influences flux into the system. In the lower panel is an example of the ‘Baseline’ configuration with integration in which bounds are modified, flux is not maintained near the reaction bound boundaries and photosystem reactions do not dictate flux into the system.

With this in mind, it is likely that the Lazy method performs better when capitalising upon light input buffering as it is less susceptible to bound changes near the photosystems, justifying the concerns associated with treating photosystem regulation similar to that of enzymatic reactions. This observation may be due to a reduction in the ability of the system to support flux near the boundaries of the upper and lower bounds, which ultimately shape the final solution. Since input flux dissipates through the system (fig. 4b), the Max configuration allows for a maintenance of flux nearer the system boundaries - which appears to be advantageous. Therefore, the dependency switch seen when using variance-guided scaling (in favour of the Max configuration) is likely due to this striking of the balance between reducing the effect of photosystem constraints early on in the system, while maintaining a larger proportion of flux near the system boundaries. It should be noted that these are speculative explanations made from mean predictions from only 7 separate datasets. Further investigation could better pinpoint key reaction bottlenecks as well as probe the influence integration methods may have deeper within the metabolic network.

### 4.0.3 Validating autotrophic metabolic models

Autotrophic constraint-based systems present unique challenges for contextualisation. The dominance of light input as an essential energy source for autotrophs means the single photon exchange reaction has a disproportionately high uptake rate compared to other essential model reactions such as bicarbonate and water exchange. Heterotrophic systems are less susceptible to the biases introduced by such a ‘top-heavy’ setup as they rely on multiple essential exchange reactions like carbohydrate, oxygen and nitrogen import - all carrying flux at rates typically within an order of magnitude of each other. In metabolic models, where all default ‘unconstrained’ reaction bounds (prior to integration) are set to the same magnitude, exchange reactions and their associated downstream pathways which support high flux rates, will be more susceptible to influence from bound modification. The subsequent disproportionate influence of these reactions (photon input in the case of autotrophically-growing organisms) on flux distributions may be viewed as problematic for valve-based integration, because only one portion of the system is likely to be modified whilst the rest of the network remains relatively unconstrained.

It is also true that an accurate light input estimate will largely dictate the demand for other exchange reactions, but the lack of influence of bound modification across multiple branches of the network may result in the failure to capture condition-specific traits of the wider system. Clearly, growth rate is intrinsically tied to energy supply from photosynthesis, meaning the ability to predict certain phenotypic traits may not be hindered by a lack of accountancy for the imbalance of autotrophic flux distributions. However, more comprehensive exploratory analyses of context-specific models would certainly benefit from these nuances being addressed.

While not directly addressing the ‘top heaviness’ of autotrophic systems, specifying bound constraints for key exchange reactions can lead to more robust solutions, with methods to quantify exchange reaction bounds having been proposed [23] [39]. Most previous validation studies, and case studies using continuous integration methods, focus on modelling cases where some experimental uptake rates are defined (even if there may be some methodological ambiguity in their determination), helping to shape final flux solutions [22] [20] [2] [38]. A given integration method’s inferred predictive performance can vary greatly, however, depending on whether these uptake rates are manually set or left unconstrained [22] [5]. While E-flux and METRADE have typically been implemented using specified uptake rates, it should also be noted that poorly substantiated input assumptions have the potential to bias outputs distributions - an important consideration since there is no uniform standard for defining such rates in autotrophic systems [23] [38]. In our analysis, we only specified the light input rate to facilitate a comparison between buffered and non-buffered configurations, leaving all other uptake rates undefined. In each case, light input flux was capped at an arbitrary baseline and subsequent growth rate predictions assessed in terms of their relative changes - an configuration similar to the ‘AC’ (all possible carbon source) setup defined in previous validation studies [20] [5] [25].

This approach was taken in the interest of maximising the number of viable conditions for analysis while also acknowledging that specific flux input rates are often not possible to obtain from previously published papers. For example, previous cyanobacterial metabolic modelling studies have calculated light available to cells using the culture surface area and dry cell weight per culture volume [39]. The light availability value can be used to set specific photon exchange reaction bounds to reflect different lighting conditions. There is, however, no strict convention for reporting the values necessary for this light availability calculation among published work. For instance, flask type, height of the culture and dry cell weight measurements (which may require inference from a calibration curve at a suitable OD [38]) are often not included in transcriptomic publications. Even in cases where absolute or relative differences in light consumption by cells can be accurately estimated, the configuration of the starting bounds should remain an important consideration, because transcript-based photosystem regulation inference may still erroneously alter the entry of flux to the wider system, even if the extent of its influence is reduced.

### 4.0.4 Limitations of the pipeline

It should be noted that full flux profiles were not analysed to infer the success of predictions, just the biomass pseudo-reaction. It could be argued therefore that the ability of transcriptomic integration to yield successful biomass predictions could be based on successful inference of only a small portion of the system. Since biomass output is closely tied to the light input to the system, success may be an indicator of this input estimation, rather than an ability to represent the whole system. The methodology presented here could be expanded to datasets which include expression profiles paired with metabolomics data, such as in Hackenberg et al. (2012) from CyanoExpress [11] - the only publication in CyanoExpress which currently contains metabolomic data. By predicting systems-level data (rather than growth rates) the pipeline could more effectively validate how well the final flux vectors represent the intracellular behaviour of the system, while also helping to quantify the extent to which growth rate predictions can serve as proxies for overall system behaviour.

Additionally, throughout this analysis, we relied upon the assumption that the true cellular objective was to maximise biomass production at every stage of growth for every condition simulated. This is because valve-based methods require the selection of an objective function in order to yield flux distributions. It should not, therefore, be overlooked that discrepancies between predictions and reference traces could be due to this oversimplification, particularly when there are alternative integration methods which do not assume biomass production as the sole cellular objective.

## 5 Conclusion

Overall, this study evaluated the impact of different transcriptomic integration methods when predicting time-series growth rates, making use of existing data from CyanoExpress. The report serves as a starting point for investigating methodological effects in cyanobacteria and proposes strategies for improving intracellular flux predictions. We identified the importance of configuration on the selection of an appropriate integration method and scaling strategy, while proposing potential justifications for these observations. We also suggest options for expanding the scope of the current pipeline by evaluating full flux distributions with the use of paired metabolomic data.

## 6 Acknowledgements

TP would like to acknowledge support of a QMUL College PhD studentship. We would also like to thank Conrad Bessant of QMUL for useful feedback on the work.

## 7 Author Contributions

TP, GH, and CDPD devised the project. TP prepared the transcriptomic and growth data sets and carried out the Flux Balance Analysis. CDPD developed the mathematical formalism for quantifying relative growth rates from OD measurements. TP drafted the manuscript and GH and CDPD helped with redrafting.

## Notes

### Competing Interest Statement

The authors have declared no competing interest.

